# Emotional contagion of pain across different social cues shares common and process-specific neural representations

**DOI:** 10.1101/2020.02.24.963595

**Authors:** Feng Zhou, Jialin Li, Weihua Zhao, Lei Xu, Xiaoxiao Zheng, Meina Fu, Shuxia Yao, Keith M. Kendrick, Tor D. Wager, Benjamin Becker

## Abstract

Insular and anterior cingulate cortex activation across vicarious pain induction procedures suggests that they are core pain empathy nodes. However, pain empathic responses encompass emotional contagion as well as unspecific arousal and overlapping functional activations are not sufficient to determine shared and process-specific neural representations. We employed multivariate pattern analyses to fMRI data acquired during physical and affective vicarious pain induction and found spatially and functionally similar cross-modality (physical versus affective) whole-brain vicarious pain-predictive patterns. Further analyses consistently identified shared neural representations in the bilateral mid-insula. Mid-insula vicarious pain patterns were not sensitive to capture non-painful arousing negative stimuli but predicted self-experienced pain during thermal stimulation, suggesting process-specific representation of emotional contagion for pain. Finally, a domain-general vicarious pain pattern which predicted vicarious as well as self-experienced pain was developed. Our findings demonstrate a generalizable neural expression of vicarious pain and suggest that the mid-insula encodes emotional contagion for pain.

## Introduction

Pain empathy, the capacity to resonate with, relate to, and share others’ pain, is an essential part of the human experience. Among other functions, it motivates helping and cooperative behaviors and aids in learning to avoid harmful situations ^1^. Vicarious pain can be elicited by multiple types of social cues, particularly the observation of an inflicted physical injury or a facial expression of pain ^2,3^. Functional magnetic resonance imaging (fMRI) studies employing corresponding experimental stimuli have identified distinct and common neural substrates pain empathy across vicarious pain induction procedures ^3^. For example, Vachon-Presseau, et al. ^4^ demonstrated that physical vicarious pain (the observation of noxious stimulation to a limp) induced greater activity in core regions of the mirror neuron system, specifically inferior frontal and posterior regions engaged in coding sensory-somatic information ^5^ while the presentation of painful facial expressions (i.e. affective vicarious pain) led to stronger increases in the medial prefrontal cortex and precuneus which have been associated with social cognitive processes such as mentalizing and theory of mind ^6–8^. Despite the different psychological domains and neural systems that mediate physical and affective vicarious pain empathy, both elicit emotional arousal and contagion for pain and commonly activate the insular and cingulate cortex, which have been suggested as shared neural representations of vicarious pain ^3,9^.

However, overlapping functional activations within these regions do not necessarily reflect shared underlying neural representations of a specific mental process ^10^, given that (1) due to local spatial dependencies the main focus of traditional mass-univariate approaches (i.e. conduct massive number of tests on brain voxels one at a time) is not on single-voxel activity, but on smoothed, regional differences in brain activity across multiple tasks or stimuli ^11^, and (2) brain regions may contain multiple, distinct populations of neurons and averaging across those neuron populations yields nonspecific signals ^10,12^. For instance, electrophysiological and optogenetic studies have identified distinct neuronal populations in the anterior cingulate and insular cortex that activate during several functional domains, including nociceptive-processing, social observation learning, empathy, reward expectancy, attention and salience ^13–19^, with mass-univariate fMRI studies suggesting that both regions are engaged by various experimental paradigms including not only experienced and observed pain, but also reward, arousal, salience and attention ^20–24^. Despite the overlapping fMRI activity in response to different experimental manipulations the underlying brain representations may be separable ^25–27^, emphasizing that more fine-grained analyses are required to determine process-specific shared or distinct neural representations ^10^.

In an effort to overcome these limitations recent studies have proposed that multivariate pattern analysis (MVPA), which can be effective in extracting information at much finer spatial scales (i.e., below the intrinsic resolution determined by the voxel size) by pooling together weak feature-selective signals in each voxel ^28,29^ and represents a more suitable approach to support or reject claims about neural mechanisms that are shared between mental processes ^10,30,31^. Multivariate pattern analysis is a machine learning technique that uses distributed patterns of voxel-wise activity to predict the neural representation of mental processes and may detect differences in the spatial distribution of fMRI signals even when single voxel activity does not differentiate between conditions ^28^. In support of this view, a growing number of recent studies have demonstrated functional independence of overlapping univariate activation in these brain regions using MVPA ^25–27,32^, including separable neural representations of physical and social rejection pain within the dorsal anterior cingulate cortex ^27^ and of modality-specific aversive experience in the anterior insular cortex ^25,26^.

Nevertheless, shared multivariate patterns do not necessarily imply process-specific common neural representations per se given that the shared neural representations could simply reflect common demands on basal processing domains such as attention or arousal ^25,26^. For instance, Corradi-Dell’Acqua, et al. ^33^ demonstrated that common local patterns between experienced and observed pain in the anterior insula were not specific to pain, but rather involve unspecific processes related to aversive emotional experience such as emotional arousal. Emotional contagion and emotional arousal are two key component of perception of other’s affective states ^34^ leading to the question of which specific component of pain empathy is shared in the common neural representations. Therefore, we propose that if the physical and affective vicarious pain-predictive patterns share functionally specific neural representations of emotional contagion for pain they should meet four criteria. Specifically, the signatures should be: (1) sensitive to within-modality vicarious pain experience; (2) exhibit similar spatial patterns which are generalizable across different vicarious pain induction procedures; (3) pain contagion specific (i.e. not sensitive to emotional arousal), and (4) able to predict the levels of self-experienced pain.

Against this background the present study employed MVPA to investigate shared global (whole-brain) and regional neural representations of vicarious pain across different modalities in healthy participants who underwent induction of physical and affective vicarious pain (i.e. stimuli showing noxious manipulations to limbs and painful facial expressions, respectively) as well as corresponding non-pain control stimuli during fMRI acquisition (**Fig. 1A**). Given that small sample size may lead to a large cross-validation error which is the discrepancy between the prediction accuracy measured by cross-validation and the expected accuracy on new data ^35^ and fMRI-based inferences on regions that are most predictive substantially benefit from larger samples ^36^ the study included a comparably large sample of n = 238 individuals [details see Methods and refs. ^37,38^]. To further test the specificity of the shared neural representations with respect to the emotional contagion of pain rather than emotional arousal the participants additionally underwent an emotion processing paradigm with neutral, positive and negative stimuli from the International Affective Picture System (IAPS) (details see Methods). Moreover, we included an independent fMRI dataset that collected ratings of self-experienced pain during thermal pain induction task details are provided in Methods and refs. ^39,40^ to determine whether the neural representations of vicarious pain could predict the level of self-experienced somatic pain.

**Fig. 1.**
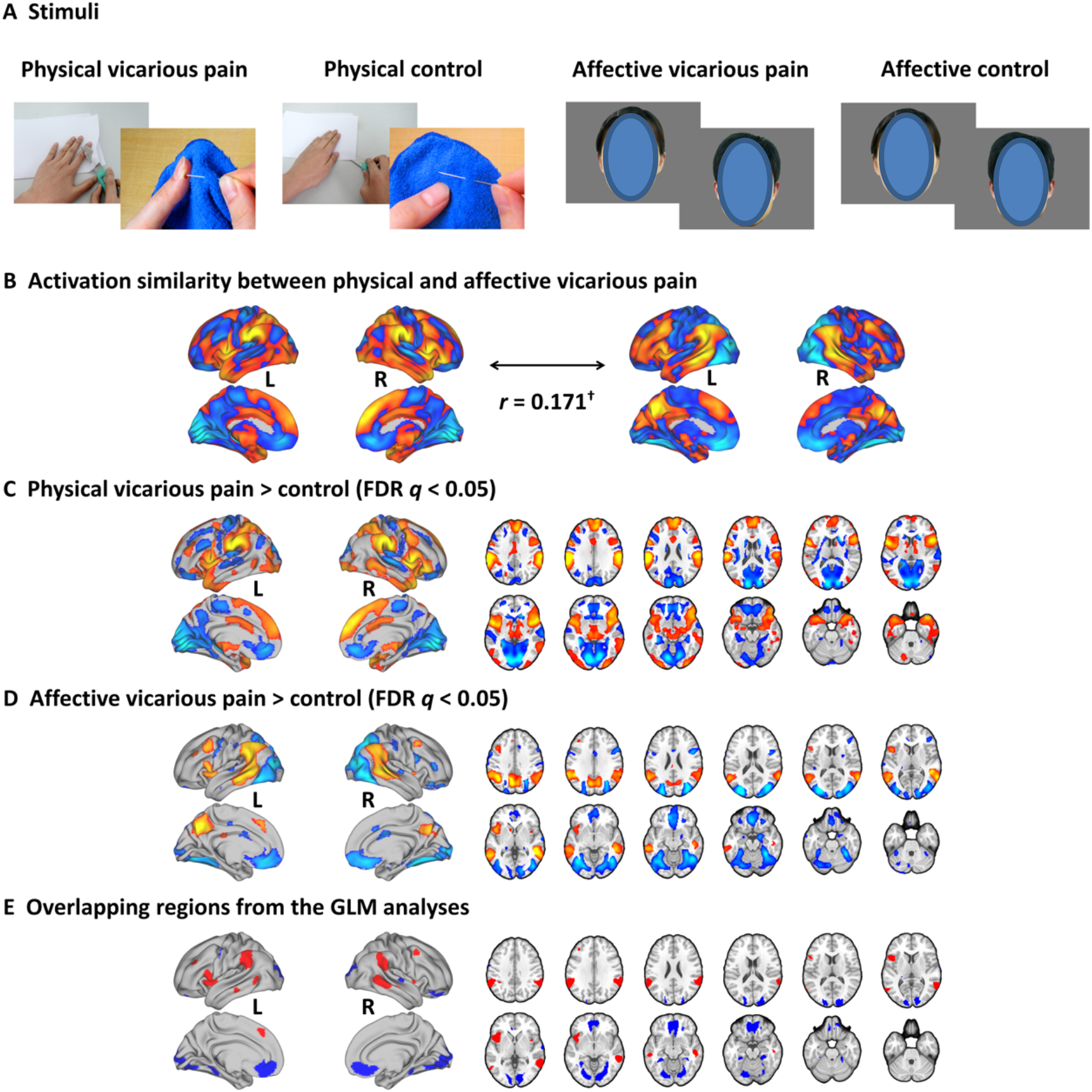
Experimental stimuli and mass-univariate analyses. (A) Examples of stimuli for physical and affective vicarious pain as well as corresponding control. For the preprint version the faces were masked with blue ovals according to recommendations of the preprint repository. (B) Physical vicarious pain activation pattern was spatially (marginally) correlated to affective vicarious pain. (C) Univariate analysis result for physical vicarious pain against physical control thresholded at FDR *q* < 0.05 (two-tailed). (D) Univariate analysis result for affective vicarious pain against affective control thresholded at FDR *q* < 0.05 (two-tailed). (E) Overlapping activation between physical and affective vicarious pain.

Through a series of analyses, we demonstrate similar (spatially correlated) whole-brain multivariate patterns across physical and affective vicarious pain induction with both patterns accurately predicting within- and between-modality vicarious pain stimuli suggesting shared neural representations of vicarious pain. Whole-brain, searchlight-based and regional-focused decoding revealed a particular role of the mid-insula across modalities emphasizing an important contribution of this regions to modality-general pain empathy. Importantly, mid-insula vicarious pain-predictive patterns were not sensitive to discriminate non-painful negative stimuli, demonstrating shared vicarious pain-specific rather than general aversive or arousal representations. Of note, the small effect sizes of comparing predicted vicarious pain versus non-pain, however, demonstrated that the mid-insula was not sufficient to encode vicarious pain processing alone. Finally, we developed a general vicarious pain-predictive pattern across physical and affective signatures for inferring pain empathy-related neural representations in future studies. Our general vicarious pain-predictive pattern was strongly related to “painful” and “pain” processing according to a Neurosynth meta-analytic decoding analysis and consistent with this it could predict physical and affective vicarious pain as well as the level of self-experienced pain in an independent dataset. Taken together, our findings suggest that physical and affective vicarious pain shares similar neural representations, particularly in the mid-insula, which might be a core region specifically encoding emotional contagion for pain

## Results

### Univariate approach - shared activations for physical and affective vicarious pain

To test whether physical and affective vicarious pain shares similar activation patterns as determined by traditional mass-univariate analyses, we performed a permutation-based correlation analysis to compare the spatial similarity between the unthresholded group-level physical vicarious pain activation (physical vicarious pain > physical control) and the affective vicarious pain activation (affective vicarious pain > affective control). Activation in response to physical vicarious pain was spatially correlated with that to affective vicarious pain (r_198,551_ = 0.171, *P* < 0.1 based on permutation tests) (**Fig. 1B**). After multiple comparisons correction (FDR corrected, *q* < 0.05, two-tailed) (**Fig. 1C,D**), distributed regions of overlapping activation were identified, including a network exhibiting increased activation during both modalities encompassing the bilateral anterior and mid-insula, dorsomedial prefrontal cortex, inferior parietal lobule, middle frontal gyrus and middle temporal gyrus, as well as a network of decreased activation, including the rostral and ventral anterior cingulate cortex, ventromedial and orbitofrontal cortex, and lingual and parahippocampal gyrus (**Fig. 1E**).

### Multivariate approach – modality general vicarious pain-predictive patterns

Previous studies suggest that pain and negative emotional processes are distributed across brain regions ^25,36,39^ and that compared to whole-brain predictive model local regions explain considerably less variance in predicting these processes ^29,41^. In an initial step we therefore developed novel whole-brain patterns to decode physical and affective vicarious pain separately. The physical vicarious pain-predictive pattern yielded an average classification accuracy of 88 ± 1.5% standard error (SE), *P* < 0.001, d = 2.13; d indicates effect size in terms of Cohen’s d (accuracy = 96 ± 1.2% SE, *P* < 0.001, d = 2.17 based on a two-alternative forced-choice test) and the affective vicarious pain-predictive pattern discriminated affective vicarious pain versus affective control with 80 ± 1.8% SE accuracy, *P* < 0.001, d = 1.64 (accuracy = 88 ± 2.1% SE, *P* < 0.001, d = 1.57 based on a two-alternative forced-choice test) with a 10-fold cross-validation procedure which was repeated 10 times, yielding 10 random partitions of the original sample. Next permutation-based correlation analysis was employed to determine the similarity between the whole-brain patterns of physical and affective vicarious pain which confirmed that the modality-specific patterns were spatially correlated (r_198,551_ = 0.170, *P* < 0.001 based on permutation tests) (**Fig. 2A**). Moreover, between-modality classification showed that the physical vicarious pain-predictive pattern could reliably discriminate affective vicarious pain versus affective control with 69% accuracy (± 3.0% SE, *P* < 0.001, d = 0.65) and that the affective vicarious pain-predictive pattern could discriminate physical vicarious pain versus physical control with 78% accuracy (± 2.7% SE, *P* < 0.001, d = 1.00) based on two-alternative forced-choice tests with a repeated 10-fold cross-validation procedure. Thresholding these neural patterns at FDR *q* < 0.05 (two-tailed, bootstrap tests with 10,000 iterations) revealed that both physical and affective vicarious pain were reliably predicted by fMRI signals in the bilateral mid-insula, left putamen and left inferior parietal lobule (**Fig. 2B**), emphasizing the importance of these regions for the shared representation of physical and affective vicarious pain. Taken together, our results confirmed shared neural representations between the different vicarious pain modalities at the whole-brain level, yet the reduced between-modality prediction effect sizes as compared to within-modality prediction effect sizes (< 50%) additionally suggest distinguishable neural representations.

**Fig. 2.**
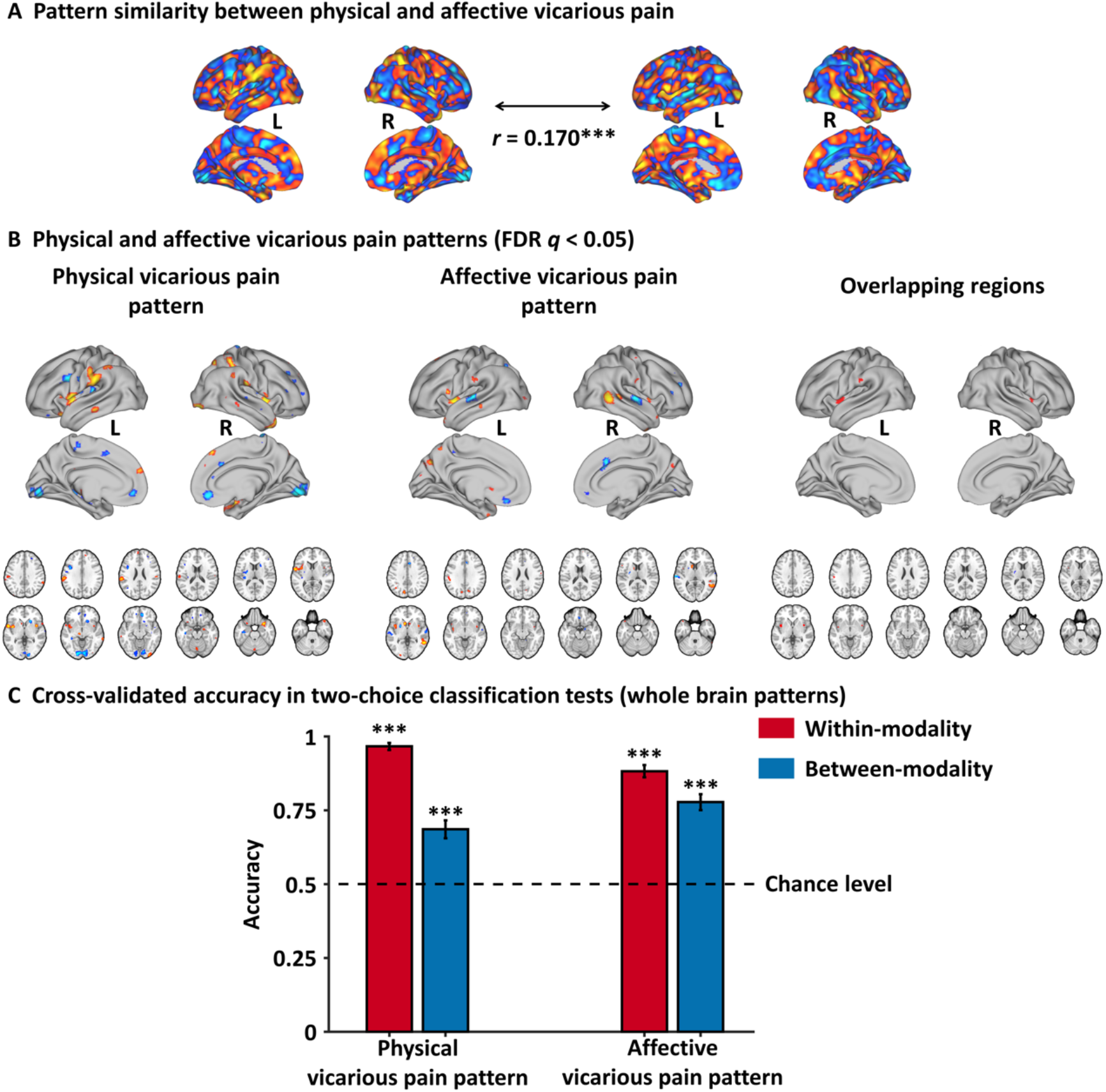
Whole-brain multivariate pattern analyses. (A) Physical vicarious pain-predictive pattern was spatially correlated to affective vicarious pain-predictive pattern. (B) Physical and affective vicarious pain-predictive patterns and overlapping reliable predictive voxels (bootstrap thresholded at FDR *q* < 0.05, two-tailed). (C) Cross-validation accuracy in two-choice classification tests based on whole-brain patterns. The results demonstrated significant within- and between- modality classifications for both physical and affective vicarious pain-predictive patterns. The dashed line indicates the chance level (50%), and the error bars represent standard error of the mean across subjects.

### Shared local representations for physical and affective vicarious pain

To identify regions with shared local patterns between physical and affective vicarious pain a within-modality cross-validation (e.g. physical vicarious pain-predictive patterns to predict physical vicarious pain versus physical control) and between-modality cross-prediction (e.g. physical vicarious pain-predictive patterns to predict affective vicarious pain versus affective control) employing a searchlight approach was conducted. A bilateral network encompassing the insula, striatum as well as the ventromedial prefrontal cortex (see **Fig. 3A**, *q* < 0.05, FDR corrected, two-tailed) demonstrated significant within-modality cross-validation and between-modality cross-prediction accuracies between physical and affective vicarious pain, implying shared representation at the local pattern level.

**Fig. 3.**
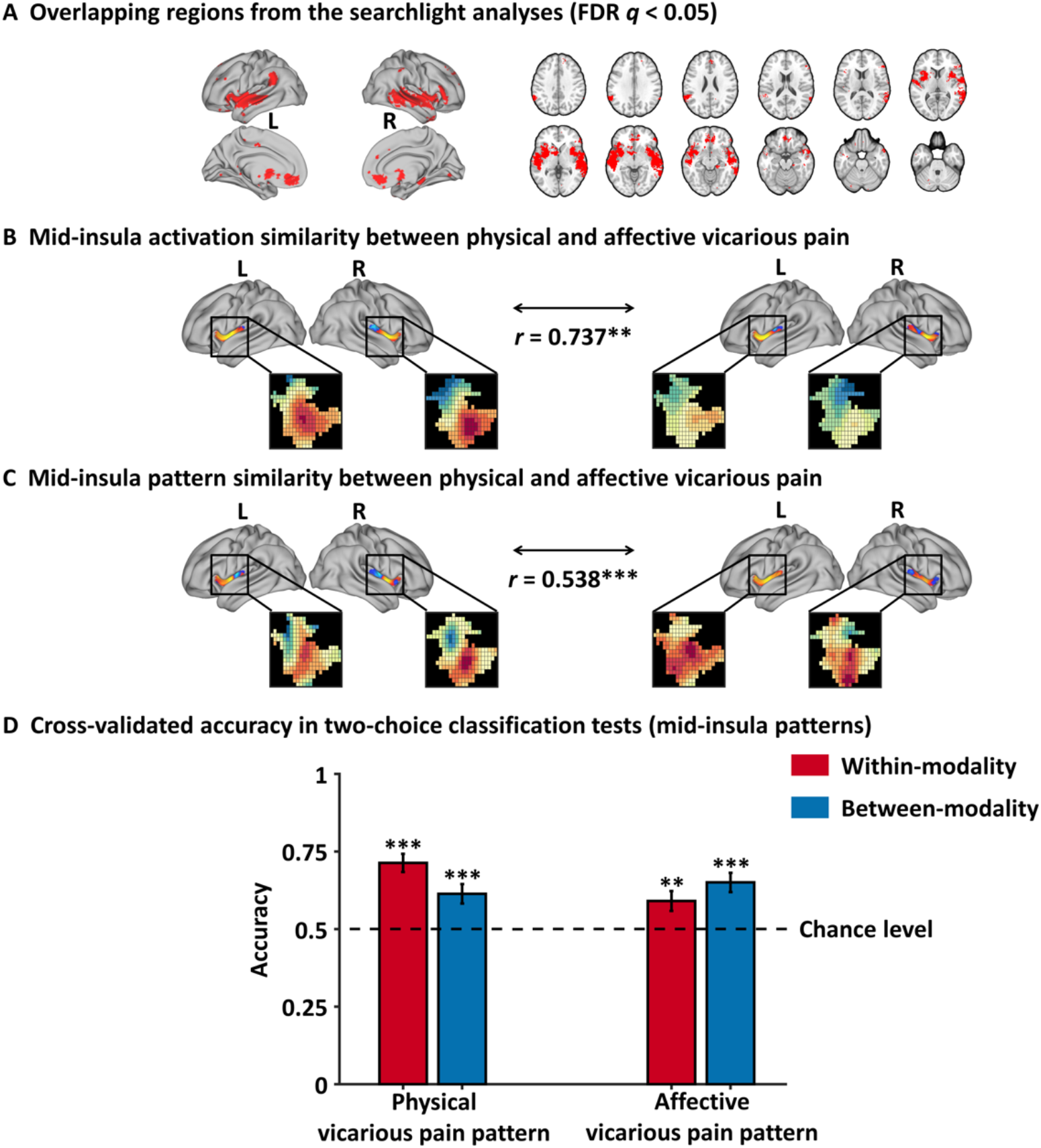
Multivariate pattern analyses with local brain regions. (A) Brain regions exhibited significant within-modality cross-validation and between-modality cross-prediction accuracies between physical and affective vicarious pain (thresholded at FDR *q* < 0.05, two-tailed). (B) Mid-insula activation to physical vicarious pain was highly similar to activation to vicarious pain. (C) Physical vicarious pain-predictive pattern in the mid-insula was spatially similar to affective pain-predictive pattern. (D) Cross-validated accuracy in two-choice classification tests with mid-insula partial patterns. The results demonstrated significant within- and between-modality classifications for both physical and affective vicarious pain-predictive patterns. The dashed line indicates the chance level (50%), and the error bars represent standard error of the mean across subjects.

### Shared representations in the mid-insula

Based on previous studies suggesting a critical role of the mid-insula in shared representations of pain across modalities (e.g. experienced and observed pain ^33^) and the present findings demonstrating overlapping activation and shared representations in the mid-insula, the similarity of the activation and pain-predictive patterns within this area (anatomically defined mid-insula from https://github.com/canlab, based on ref. ^42^) was further examined. Mid-insula focused analysis revealed that physical vicarious pain activation in the insula was strongly positively correlated with affective vicarious pain activation (r_1,947_ = 0.737, *P* = 0.006 based on permutation tests) and consistent with this, that the physical vicarious pain-predictive and affective vicarious pain-predictive pattern weights within the mid-insula were also strongly positively correlated (r_1,947_ = 0.538, *P* < 0.001 based on permutation tests) (**Fig. 3B, C**). Moreover, two-alternative forced-choice tests revealed that the mid-insula partial physical vicarious pain-predictive pattern classified above chance for both, physical vicarious pain versus physical control (71 ± 3.0% SE, *P* < 0.001, d = 0.72; within-modality) and affective vicarious pain versus affective control (61 ± 3.2% SE, *P* < 0.001, d = 0.36; between-modality prediction) in out-of-sample participants through a repeated 10-fold cross-validation procedure. In line with this, the mid-insula partial affective vicarious pain-predictive pattern discriminated physical vicarious pain versus physical control with 65% accuracy (± 3.1% SE, *P* < 0.001, d = 0.58; between-modality) and affective vicarious pain versus affective control with 60% accuracy (± 3.2% SE, *P* = 0.004, d = 0.27; within-modality) (**Fig. 3D**). Together, these findings converge on common representations of vicarious pain in the mid-insula across univariate and multivariate patterns for physical and affective vicarious pain. However, the relative low accuracies and small effect sizes compared to the prediction results with whole-brain patterns demonstrated that the mid-insula was not sufficient to capture vicarious pain processing alone.

### Shared vicarious pain representations are not sensitive to arousal or negative affect

One key question is whether the developed vicarious pain-predictive patterns are specific to emotional contagion for pain or are rather generally sensitive to emotional arousal or negative affect. To test the functional specificity whole-brain patterns were separately employed to discriminate processing of highly arousing non-painful negative from low arousing neutral stimuli from the IAPS database with two-alternative forced-choice tests through a repeated 10-fold cross-validation procedure. This approach revealed statistically significant yet comparably low accuracies and small effect sizes (physical vicarious pain-predictive pattern: 58 ± 3.2%, *P* = 0.024, d = 0.34; affective vicarious pain-predictive pattern: 61 ± 3.2% SE, *P* = 0.001, d = 0.42). In contrast testing whether shared representations in the mid-insula could discriminate negative versus neutral stimuli revealed that neither of the insula partial patterns could classify negative stimuli above chance level (physical: 56 ± 3.2% SE, *P* = 0.079, d = 0.11; affective: 56 ± 3.2% SE, *P* = 0.111, d = 0.09), suggesting a pain-specific representation in this region.

### A general across physical and affective vicarious pain-predictive pattern

Given that the physical and affective vicarious pain-predictive patterns shared similar whole-brain as well as local neural representations, we developed a general across physical and affective vicarious pain pattern which yielded a classification accuracy of 82 ± 1.2% SE, *P* < 0.001, d = 1.77 (accuracy = 91 ± 1.3% SE, *P* < 0.001, d = 1.74 based on a two-alternative forced-choice test) in discriminating vicarious pain versus non-painful control. More specifically, the pattern could accurately predict both physical vicarious pain from the physical control (95 ± 1.4% accuracy, *P* < 0.001, d = 2.10) and affective vicarious pain from the vicarious control (87 ± 2.1% accuracy, *P* < 0.001, d = 1.45), but performed considerably worse classifying non-painful negative versus neutral stimuli (59 ± 2.1% accuracy, *P* = 0.01, d = 0.30), in forced-choice classifications. In line with the spatially overlapping modality-specific vicarious pain patterns the general vicarious pain pattern was highly similar with both, the physical vicarious pain pattern (r_198,551_ = 0.587, permutated *P* < 0.001 based on permutation tests) and affective vicarious pain pattern (r_198,551_ = 0.702, *P* < 0.001 based on permutation tests). To functionally characterize the general vicarious pain-predictive pattern the Neurosynth decoder function was used to assess its similarity to the reverse inference meta-analysis maps generated for the entire set of terms included in the Neurosynth dataset. The most relevant features were “painful” and “pain” [for the top 50 terms (excluding anatomical terms) ranked by the correlation strengths between the vicarious pain pattern and the meta-analytic maps see word cloud (the size of the font was scaled by correlation strength, **Fig. 4A**). After thresholding and correction for multiple comparisons (bootstrapping 10,000 samples, FDR q < 0.05, two-tailed), the general vicarious pain-predictive pattern revealed a distributed network engaged in vicarious pain processing encompassing the bilateral mid-insula, inferior parietal lobule and ventromedial prefrontal cortex (**Fig. 4B**), which suggested the importance of these regions in encoding vicarious pain.

**Fig. 4.**
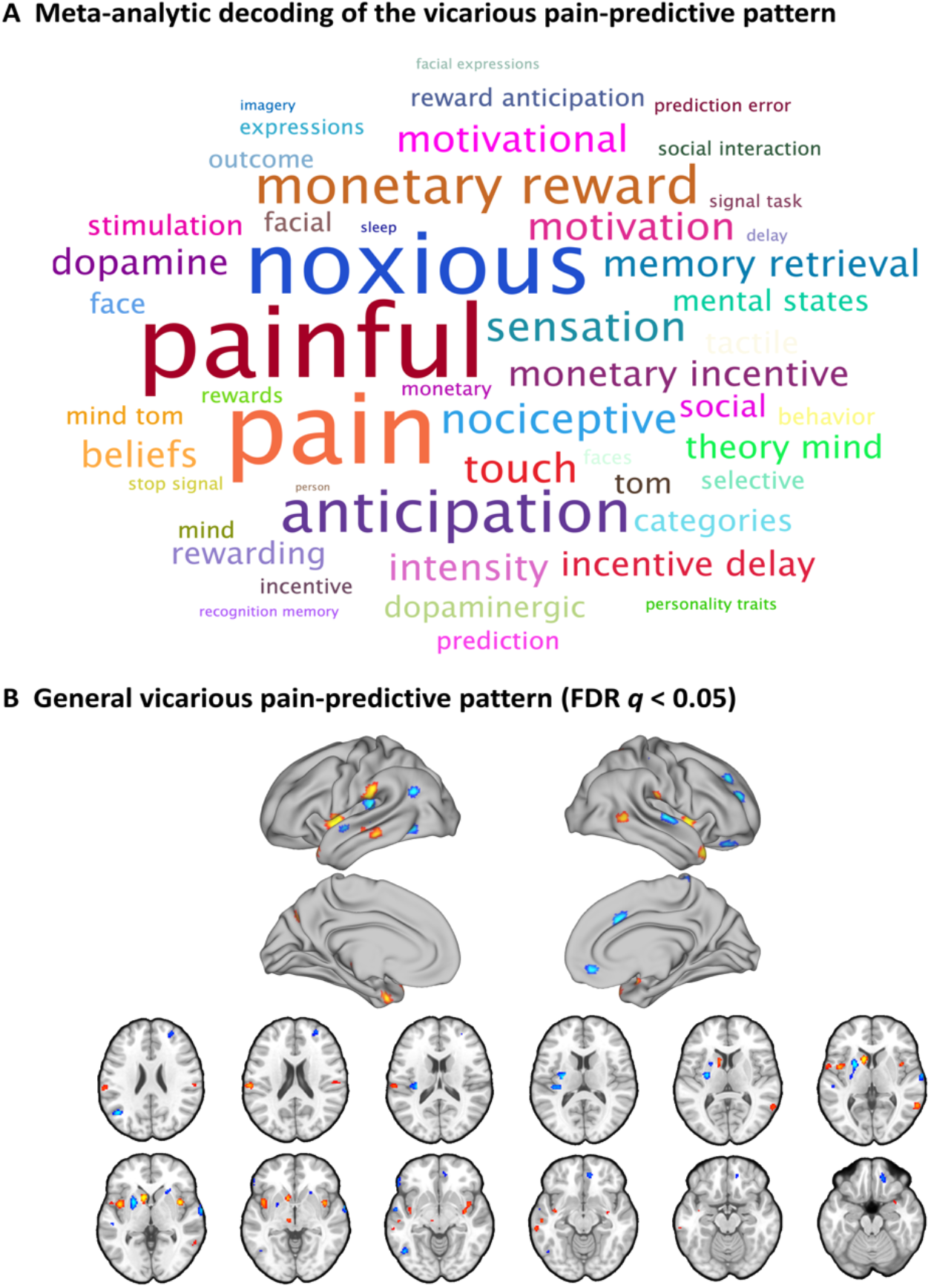
A general across physical and affective vicarious pain-predictive pattern. (A) Word cloud showing the top 50 relevant terms (excluding anatomical terms) for the meta-analytic decoding of the general vicarious pain-predictive pattern. The size of the font was scaled by correlation strength. (B) When thresholded at FDR *q* < 0.05, two-tailed (bootstrapped 10,000 samples) the general vicarious pain-predictive pattern revealed a distributed network of vicarious pain empathy representation including bilateral mid-insula and ventromedial prefrontal cortex.

### Generalization and functional relevance of the vicarious pain-predictive pattern

To test the generalizability of the identified neural representations of general vicarious pain, we applied the whole-brain general vicarious pain-predictive pattern to self-experienced thermal pain data using dot-product of vectorized activation maps with the pattern classifier weights. We found that the general vicarious pain-predictive pattern expressions were highly correlated with both overall objective temperature levels (r_196_ = 0.538, *P* < 0.001) and subjective pain ratings (r_196_ = 0.507, *P* < 0.001). Moreover, the general pain-predictive pattern discriminated high thermal pain versus low thermal pain with a 94% accuracy (± 4.2% SE, *P* < 0.001, d = 2.00), high thermal pain versus medium thermal pain with a 91% accuracy (± 5.0% SE, *P* < 0.001, d = 1.56) and medium thermal pain versus low thermal pain with an 82% accuracy (± 6.7% SE, *P* = 0.001, d = 1.20) using two-alternative forced-choice tests (**Fig. 5A**).

**Fig 5.**
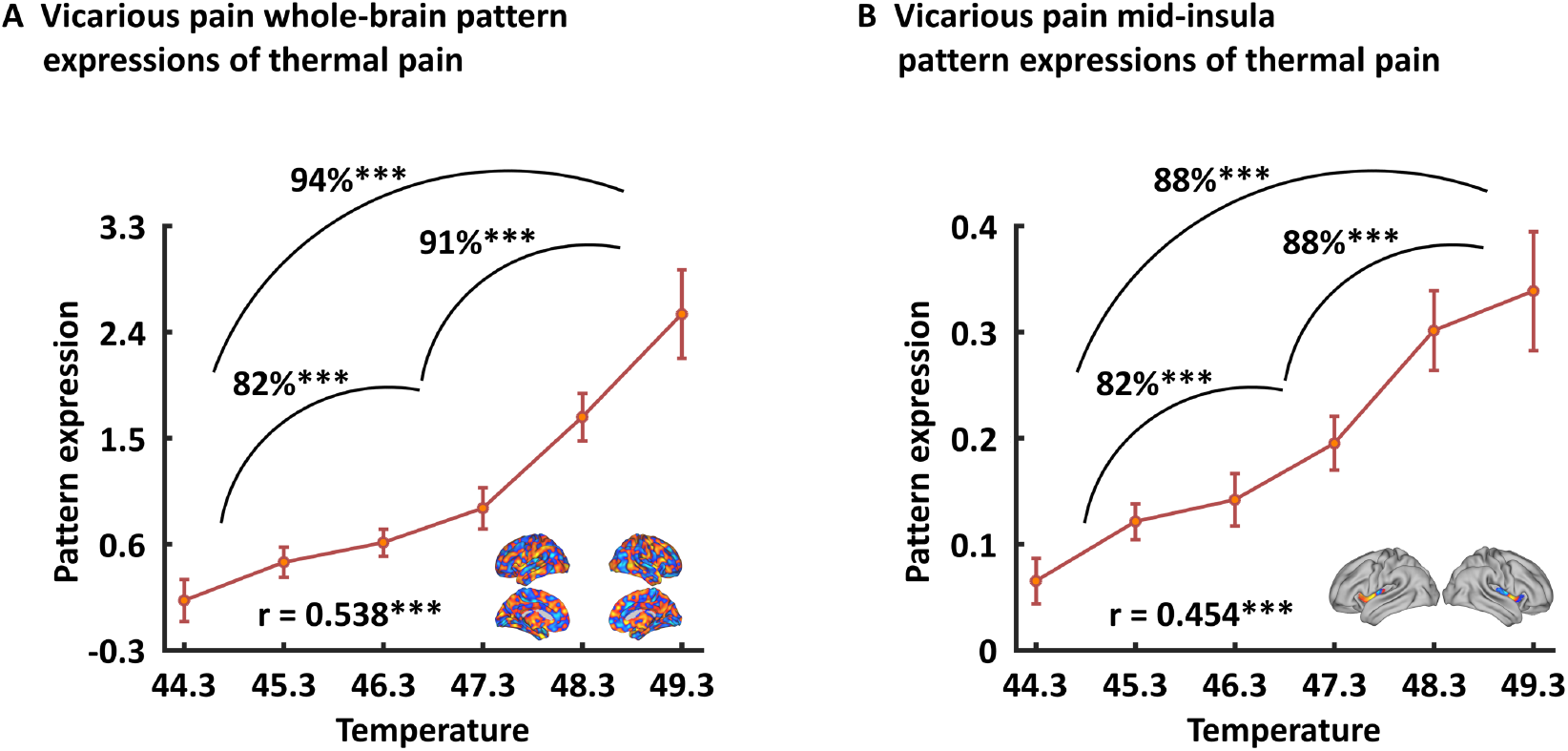
Generalizability of the general across physical and affective vicarious pain-predictive pattern. Both whole-brain (A) and mid-insula (B) vicarious pain-predictive patterns could accurately predict the severity and classify the levels of self-experienced pain in an independent dataset.

When prediction focused on the mid-insula the general vicarious pain-predictive local pattern could discriminate high thermal pain versus low thermal pain (accuracy = 88 ± 5.7% SE, *P* < 0.001, d = 1.56), high thermal pain versus medium thermal pain (accuracy = 88 ± 5.7% SE, *P* < 0.001, d = 1.23) and medium thermal pain versus low thermal pain (accuracy = 82 ± 6.7% SE, *P* < 0.001, d = 1.49) above chance levels (**Fig. 5B**). In addition, the mid-insula partial pattern expressions (i.e. focusing on the mid-insula pattern) were highly correlated with temperature levels (*r*_196_ = 0.454, *P* < 0.001) as well as individual pain ratings (*r*_196_ = 0.440, *P* < 0.001). Together with the predictions using physical and affective vicarious pain-predictive patterns separately (see **Supplementary Results** for details), our results demonstrated that the vicarious pain patterns shared a common neural basis (especially in the mid-insula) which might be linked to the affective experience of pain, specifically the strengths of experienced pain including physically-induced pain.

## Discussion

Several studies have explored the neural underpinnings of vicarious pain in humans and suggested overlapping univariate fMRI activity in the anterior cingulate and insular cortex across different vicarious pain induction procedures (for meta-analyses see e.g. refs. ^3,9^). However, the conventional univariate approach lacks anatomical and functional specificity to test the question of whether vicarious pain across different modalities shares common and process-specific neural representations ^10,25,27,43^. By using fine-grained MVPA, which is sensitive and specific to particular types of mental processes including pain ^29,32,39^ and may reflect population codes across neurons ^44,45^, we demonstrated similar (i.e. spatially correlated) multivariate patterns encoding physical and affective vicarious pain at the whole-brain level and within the mid-insular cortex. Furthermore, we demonstrated that these patterns were not sensitive to infer the processing of non-painful negative arousing stimuli in the same sample but could predict the objective and subjective intensity of self-experienced thermal pain in an independent sample. Together these results provided evidence for generalized neural representation of vicarious pain, particularly in the mid-insula, and demonstrated that the shared neural signature may specifically capture the emotional contagion of pain that characterizes vicarious pain experience.

The idea that vicarious pain across different induction procedures shares common neural representations has been supported by meta-analyses covering previous fMRI pain empathy studies that demonstrated overlapping activation in the insular and cingulate cortex ^3,9^. Consistent with these meta-analytic findings we found that these regions were consistently engaged during both physical and affective vicarious pain. However, the insular and anterior cingulate cortex are involved in a wide range of mental processes including representation of interoceptive and affective states as well as salience detection ^9,22,46,47^, suggesting that the overlapping activity might be due to common underlying mental processes such as detecting and orienting attention towards salient stimuli or unspecific emotional arousal ^26,33,48^. Thus, we employed a more fine-grained and well-controlled analysis which may determine specific representations of vicarious pain contagion in meso-scale neural circuits were.

To systematically test whether emotional contagion for pain elicited by different social cues shares common neural representations we developed and compared multivariate patterns that predicted physical and affective vicarious pain stimuli respectively. While mass-univariate analysis results reflect the presence of intermingled neuronal populations related to stimulus-specific representations, MVPA investigates whether idiosyncratic spatial variations in the fMRI signal are shared or dissociated across different conditions and thus might be more suitable to determine process-specific representations in meso-scale neural circuits ^28,30,45^. Moreover, previous studies suggest that whole-brain predictive models could better capture emotional processes compared to regional approaches, such as decoding of a single brain region or searchlight-based methods ^29,41^.To this end, we first identified whole-brain fMRI patterns that accurately predicted physical and affective vicarious pain, respectively. We found that the physical and affective vicarious pain-predictive patterns were spatially correlated and both could classify within- and between-modality painful versus non-painful stimuli at the whole-brain level, suggesting that physical and affective vicarious pain shares distributed processing across multiple systems and component processes. Notably, decreased accuracies and effect sizes in the cross-modality additionally suggest distinguishable neural representations of vicarious affective and physical reflecting the engagement of different component processes.

In the context of previous studies suggesting that pain empathy deficits are mediated by regional-specific brain lesions and functional dysregulations ^49–51^ the question for the contribution of specific brain regions arises. Thresholding the vicarious pain patterns (at FDR *q* < 0.05, two-tailed) allowed us to identify voxels that reliably contributed to the respective decoders and revealed that specifically the bilateral mid-insula provided important features to predict both physical and affective vicarious pain. Moreover, the mid-insula partial vicarious pain patterns were highly spatially correlated and both could significantly predict within- and between-modality vicarious pain-related experience. Consistent with this, searchlight-based classification analyses also demonstrated that mid-insula local patterns produced significant within- and between-modality predictions of vicarious pain. Our results are in line with a previous meta-analysis showing that the mid-insula responds specifically to empathy for pain across different task paradigms compared to empathy for non-pain negative affective states ^9^, which together with the present findings suggests that the mid-insula represents a core neural substrate for vicarious pain. Although multivariate predictive models can capture information at much finer spatial scales and consequently anatomical specificity ^28,29^ the question of the specific mental processes captured by our vicarious pain predictive patterns remains unclear. Pain empathy is a multi-component process that includes emotional contagion but also emotional arousal and negative affect and these unspecific processes may be captured by the decoders. To determine the functional specificity of the neural representations we applied the vicarious pain-predictive patterns to data from an emotion processing paradigm acquired in the same sample as well as to data from a thermal pain induction experiment in an independent sample and found that (1) the vicarious pain patterns performed only modest for discriminating neural representations of high arousing (non-painful) negative stimuli from low arousing neutral stimuli, and (2) the whole-brain but specifically the mid-insula patterns predicted levels of self-experienced thermal pain with high accuracy. Finally, we developed a general vicarious pain-predictive pattern across physical and affective vicarious pain induction procedures and demonstrated that it accurately predicted both physical and affective vicarious pain (accuracies > 87%) as well as thermal pain intensities (accuracies > 82%), yet classified non-painful negative versus neutral stimuli with comparably low accuracy (59%). In line with the prediction results, meta-analytic decoding analysis revealed that this general vicarious pain pattern was highly correlated with the domains of “painful” and “pain”, but not with “arousal”, “valence” or “negative” (not shown in the top 100 relevant terms). Together these findings suggest a shared neural representation of vicarious pain and a high-specificity of the whole-brain and specifically the mid-insula patterns for the emotional contagion component of the pain empathic response. A previous study developed a vicarious pain signature (VPS) that was sensitive and specific to physical vicarious pain, but not sensitive to the intensity of self-experienced somatic pain ^25^. Examining similarities with our general vicarious pain-predictive pattern revealed only modest spatial correlations between the two patterns (r = 0.04). The different instructions employed in the experiments might have contributed to the low overlap such that participants in the previous study were required to explicitly rate their emotional response to the stimuli whereas we decided for a passive viewing paradigm to control for cognitive processes which can modulate emotional contagion and painful experience as well as the specific neural networks engaged ^3,52^. Moreover, the present pattern could successfully predict pain experience during thermal heat stimulation while the VPS was not sensitive to self-experienced pain, suggesting that the passive presentation may have better captured the emotional contagion component of vicarious pain.

Our results highlighted the mid-insula as a key region sharing similar neural representations across physical and affective vicarious pain suggesting that it may contribute to the core emotional contagion component that characterizes vicarious pain experience. Consistent with whole-brain results, the shared information in the mid-insula was specific to vicarious pain rather than negative affect or arousal. Previous non-human primate and human studies indicate that the posterior and mid-insula receive nociceptive information from thalamic nuclei ^53,54^ which are in turn conveyed to the anterior insula for progressive integration with higher level affective and interoceptive experience ^26,55^. In line with these findings, previous studies reported engagement of the anterior insula in response to a range of affective and salient stimuli while the mid-insula seems to be more specifically involved in pain-related experiences ^9,33^. Studies examining the functional and anatomical organization of the insular cortex with intracerebral electrical stimulation have demonstrated that painful sensations are elicited by stimulation of the mid-insula ^56^. Taken together, the functional relevance of the mid-insula to predict objective and subjective pain experience in an independent sample and the contribution of this region to nociception as well as vicarious pain ^9,57,58^ suggests that the shared representations across vicarious pain induction procedures may specifically code the contagious experience of pain. However, consistent with previous evidence that (physical) vicarious pain representation is distributed across brain regions and single local regions exhibit considerably lower effect sizes compared to whole-brain predictive models ^25^ we found that the prediction effect sizes for the mid-insula were smaller than those observed in our whole brain analyses. These findings suggest that despite the key role of the mid-insula in pain contagion this region is not sufficient to fully capture this process. Notably, although the vicarious pain patterns could accurately predict the intensity of self-experienced pain this result does not imply that vicarious and somatic pain share common neural representations (for more detailed investigation see e.g. ref. ^25^).

Consistent with previous studies suggesting that the anterior cingulate cortex represents a core brain region for emotional empathy in general and pain empathy in particular ^3,9,59^, we found overlapping deactivations in the rostral and ventral anterior cingulate cortex in mass-univariate analyses and shared patterns in the dorsal, rostral and ventral anterior cingulate cortex in searchlight-based prediction analyses between physical and affective vicarious pain. However, no overlapping reliable predictive voxels for whole-brain physical and affective pain-predictive patterns were found in cingulate regions suggesting a differential involvement of this region during affective and physical vicarious pain induction procedures. From a methodological perspective these results may reflect that the whole-brain predictive model could provide a more specific neural description of a behavior or mental process ^29,41^. In line with our findings previous studies also show that significant activation and searchlight-based prediction in local regions do not necessarily imply reliable predictive features in whole-brain predictive models ^25,27^. From a brain systems perspective these findings may indicate that the anterior cingulate cortex is not specifically involved in the contagious pain empathic response elicited across induction procedures. Although the anterior cingulate cortex has been reliably identified in meta-analytic studies covering brain activation patterns during (pain) empathy induction procedures (see e.g. refs. ^3,9^) it has also been associated with a number of basal processes, including arousal and salience, and activation in this region may reflect rather unspecific neural responses. Although post-scanning ratings confirmed that – relative to their corresponding control stimuli – physical and affective pain stimuli were perceived as more painful they were also perceived as more arousing (all *P*s < 0.001, details see **Supplementary Table 1**) the additional inclusion of the IAPS task allowed us to exclude unspecific contributions of arousal related neural activation.

In conclusion, by applying a novel whole-brain as well as local-region based MVPA approaches in a large sample of healthy adults, our results provide the first neuroimaging evidence that physical and affective vicarious pain shares neural representations, especially in the mid-insula which may specifically encode the emotional contagion component of vicarious pain experience. Moreover, we also provide a general vicarious pain-predictive pattern (across physical and affective vicarious pain stimuli), which may be employed in future studies to facilitate inferences about pain empathy across modalities as well as self-experienced pain. Our study offers a new approach to better understand pain empathy by exploring common neural representations and linking these shared representations to felt pain.

## Methods

### Participants

N = 252 healthy young participants were enrolled in the current study and underwent a previously validated physical and affective vicarious pain empathy fMRI paradigm. The fMRI data on the basic group activation maps for physical and affective vicarious pain contrasts were previously published in a study examining dimensional associations with trait autism and alexithymia ^37^ and a study investigating network-level communication during pain empathic processing using an exploratory inter-subject phase synchronization approach ^38^. Of note, the aim, methodological approach and hypotheses of the current study were independent from these previous publications; here we focus on identifying an fMRI multivariate pattern for physical and affective vicarious pain separately and assessing their relationship. To further examine the specificity of the determined pain patterns from general negative emotion processing the data from an emotion processing paradigm from the same subjects was additionally used. Due to technical issues during data acquisition (incomplete data, n = 6), left-handedness (n = 4) or excessive head motion (> 3 mm translation or 3° rotation; n = 4) data from 14 participants were excluded leading to a sample of n = 238 participants (118 females; mean ± SD age = 21.58 ± 2.32 years) for the pain empathy analyses; data from 15 participants (incomplete data n = 8; left-handedness, n = 4; excessive head motion, n = 3) was excluded from the emotion processing paradigm analyses (n = 237; 120 females; mean ± SD age = 21.55 ± 2.30 years). All participants provided written informed consent, the study was approved by the local ethics committee at the University of Electronic Science and Technology of China and was in accordance with the most recent revision of the Declaration of Helsinki.

### Pain empathy paradigm

The pain empathy paradigm employed a blocked design incorporating previously validated visual stimuli displaying painful everyday scenes (physical vicarious pain) and painful facial expressions (affective vicarious pain) as well as modality-specific control stimuli displaying non-painful scenes (physical control) or neutral facial expressions (affective control), respectively ^60–62^. The physical vicarious stimuli displayed a person’s hand or foot in painful or non-painful everyday situations from the first-person perspective (e.g. the painful stimulus displays cutting a hand with a knife whereas the matched non-painful control stimulus shows cutting vegetables with a knife). The affective vicarious stimuli incorporated painful and neutral facial expressions from 16 Chinese actors (8 males) (see **Fig. 1A** for examples). A total of 16 blocks (4 blocks per condition) were presented in a pseudo-randomized order and interspersed by a jittered red fixation cross (8, 10, or 12s). Each block (16s) incorporated four condition-specific stimuli (each presented for 3s) separated by a white fixation cross (1s).

### Emotion processing paradigm

In line with the pain empathy paradigm the emotion processing paradigm employed a block design incorporating three experimental conditions (positive, negative and neutral pictures). All stimuli were from the International Affective Picture System (IAPS) database. A total of 19 blocks (neutral, 7 blocks; negative, 6 blocks; positive, 6 blocks) were presented in a pseudo-randomized order and interspersed by a jittered red fixation cross (8, 10, or 12s). In each block (16s) four condition-specific stimuli (3s) were presented and separated by a white fixation cross (1s). Participants were asked to press a button when a stimulus with a white frame (one in each block) was presented. We recruited an independent sample of 37 subjects (16 females; Mean ± SD age =23.60 ± 2.86 years) to rate the valence and arousal for each stimulus with nine-point Likert scales (1 = “very negative” or “lowly arousing”, 9 = “very positive” or “highly arousing”). Negative stimuli elicited substantial negative affect and arousal on numerical rating scales (mean ± SD valence = 2.41 ± 0.96; mean ± SD arousal = 6.34 ± 1.34) compared with neutral stimuli (mean ± SD valence = 5.35 ± 0.50; mean ± SD arousal = 3.22 ± 1.54; t_36_ = −16.09, *P* < 0.001; t_36_ = 12.65, *P* < 0.001, respectively).

### MRI data acquisition and preprocessing

MRI data were collected on a 3.0-T GE Discovery MR750 system (General Electric Medical System, Milwaukee, WI, USA). Functional MRI data was acquired using a T2*-weighted echo-planar imaging (EPI) pulse sequence (repetition time = 2s, echo time = 30ms, 39 slices, slice thickness = 3.4mm, gap = 0.6mm, field of view = 240 × 240mm, resolution = 64 × 64, flip angle = 90°, voxel size = 3.75 × 3.75 × 4mm). To improve spatial normalization and exclude participants with apparent brain pathologies a high-resolution T1-weighted image was acquired using a 3D spoiled gradient recalled (SPGR) sequence (repetition time = 6ms, echo time = minimum, 156 slices, slice thickness = 1mm, no gap, field of view = 256 × 256mm, acquisition matrix = 256 × 256, flip angle = 9°, voxel size = 1 × 1 × 1mm). OptoActive MRI headphones (http://www.optoacoustics.com/) were used to reduce acoustic noise exposure for the participants during MRI data acquisition.

Functional MRI data was preprocessed using Statistical Parametric Mapping (SPM12, https://www.fil.ion.ucl.ac.uk/spm/software/spm12/). The first 10 volumes of each run were discarded to allow MRI T1 equilibration and active noise cancelling by the headphones. The remaining volumes were spatially realigned to the first volume and unwarped to correct for nonlinear distortions related to head motion or magnetic field inhomogeneity. The anatomical image was segmented into grey matter, white matter, cerebrospinal fluid, bone, fat and air by registering tissue types to tissue probability maps. Next, the skull-stripped and bias corrected structural image was generated and the functional images were co-registered to this image. The functional images were subsequently normalized the Montreal Neurological Institute (MNI) space (interpolated to 2 × 2 × 2mm voxel size) by applying the forward deformation parameters that were obtained from the segmentation procedure, and spatially smoothed using an 8-mm full-width at half maximum (FWHM) Gaussian kernel.

### Pain empathy – univariate general linear model (GLM) analyses

A two-level random effects GLM analysis was conducted on the fMRI signal to determine shared modality-specific activation patterns using a mass-univariate GLM approach. The first-level model included four condition-specific (physical vicarious pain, physical control, affective vicarious pain, and affective control) box-car regressors logged to the first stimulus presentation per block that were convolved with SPM12’s canonical hemodynamic response function (HRF). The fixation cross epoch during the inter-block interval served as implicit baseline, and a high-pass filter of 128 seconds was applied to remove low frequency drifts. Regressors of non-interest (nuisance variables) included (1) six head movement parameters and their squares, their derivatives and squared derivatives (leading to 24 motion-related nuisance regressors in total); and (2) motion and signal-intensity outliers (based on Nipype’s rapidart function). Single-subject voxel-wise statistical parametric maps for the empathy modality-specific contrasts (physical vicarious pain > physical control and affective vicarious pain > affective control) were obtained and subjected to group-level one-sample t-tests. The corresponding analyses were thresholded and corrected for multiple comparisons within a grey matter mask based on false discovery rate (FDR *q* < 0.05, two-tailed) with a minimum extent of 100mm^3^. The resulting thresholded activation maps were next used to identify common regions of activation across the modalities (physical and affective vicarious pain; i.e. masking the overlapping significant voxels).

To determine the activation similarity of physical and affective vicarious pain a permutation-based correlation analysis was employed ^63^. Specifically, we (1) calculated Pearson’s correlation (r) between the modality-specific unthresholded statistical maps (physical vicarious pain > physical control versus affective vicarious pain > affective control), (2) shuffled the condition labels for the physical stimuli, obtained a new group-level statistical map for “physical vicarious pain > physical control” and calculated the activation similarity of affective and the “modelled” physical vicarious pain, (3) repeated step (2) 10,000 times, (4) repeated steps (2-3) with shuffled labels for affective instead of physical stimuli, and finally (5) calculate the probability of observing the activation similarity between the true physical and affective pain given the null distribution of permuted activation similarity. A *P* value < 0.05 was being considered statistically significant and between 0.05 - 0.1 was being considered as marginal significant.

### Pain empathy - multivariate pattern analyses

For the multivariate pattern analyses, nuisance regression (24 head motion parameters, motion and signal-intensity outliers, and linear trend) and high-pass filtering (cut off at 128s) were initially simultaneous conducted on the preprocessed fMRI data. Next, the fMRI signal was averaged within the 4 condition-specific blocks (shifted by 3 TRs to account for the delay of the HRF). Linear support vector machines (SVMs, C = 1) were then employed to the whole-brain maps (restrict to a grey matter mask) to train multivariate pattern classifiers on the cleaned averaged fMRI signal to discriminate physical vicarious pain versus physical control and affective vicarious pain versus affective control separately. The classification performance was evaluated by a 10-fold cross-validation procedure during which all participants were randomly assigned to 10 subsamples of 23 or 24 participants using MATLAB’s cvpartition function. The optimal hyperplane was computed based on the multivariate pattern of 214 or 215 participants (training set) and evaluated by the excluded 24 or 23 participants (test set). The training set was linearly scaled to [−1, 1], and the test set was next scaled using the same scaling parameters before applying SVM ^64^. This procedure was repeated 10 times with each subsample being the testing set once. To avoid a potential bias of training-test splits, the cross-validation procedures throughout the study were repeated 10 times by producing different splits in each repetition and the resultant accuracy and *P* values were averaged to produce a convergent estimation ^65^. In line with the mass-univariate analyses and to identify which brain regions made reliable contributions to the decoders ^39,66^, the pattern maps were thresholded at FDR *q* < 0.05 (two-tailed) with a minimum extent of 100mm^3^ using bootstrap procedures with 10,000 samples. Next the thresholded maps were subjected to a conjunction analysis to identify regions that robustly contributed to both physical and affective vicarious pain classifiers by masking overlapping significant voxels. Statistical maps were visualized using the Connectome Workbench provided by the Human Connectome Project (https://www.humanconnectome.org/software/connectome-workbench).

Similarity patterns between the modality-specific neural patterns were determined employing (1) Pearson’s correlation between the whole-brain unthresholded classifier weights using a permutation test (similar to the activation similarity analysis), and (2) “between - modality classification” tests encompassing the following two steps: (a) pattern classifiers were trained separately for physical vicarious pain versus physical control and affective vicarious pain versus affective control with a 10-fold cross-validation procedure (repeated 10 times), and next (b) applying the identified patterns of physical and affective vicarious pain to out-of-sample participants for the affective vicarious pain versus affective control and physical vicarious pain versus physical control respectively using a two-alternative forced choice test, where pattern expression values were compared for two conditions with the image exhibiting the higher expression being determined as pain.

### Pain empathy – within- and between- modality classification analyses employing local classifiers

To further identify regions with shared neural expressions across physical and affective vicarious pain, a local pattern-based classification approach with three-voxel radius spherical searchlights around center voxels was employed ^27,33^. Specifically, (1) multivariate pattern classifiers using a defined local region were trained to discriminate vicarious pain versus control within each modality (i.e. physical and affective stimuli) separately, (2) the patterns obtained were next applied to out-of-sample participants for within-modality cross-validation and between-modality cross-prediction. Steps (1) and (2) were repeated for each local region across the whole-brain. It was hypothesized that shared neural representations for physical and affective pain within a local region would be reflected by significant cross-validation and cross-prediction accuracies for each classifier.

### Specificity of the physical and affective vicarious pain-predictive patterns

To test whether the observed physical and affective vicarious pain-predictive patterns were specific to pain processing or rather reflect general aspects of negative emotional processing the two pain predictive patterns were applied to the data from the emotional task paradigm. The first-level model for the emotion processing data included the four experimental conditions (positive, negative, neutral and white framed stimuli) and high-pass filter and nuisance regressors were identical to the pain empathy GLM analysis. The two pain-predictive patterns were next applied to negative and neutral contrasts (via dot-products) using a repeated 10-fold cross-validation procedure separately, and subsequently two-alternative forced choice tests were employed to discriminate negative versus neutral stimuli.

### Generalized vicarious pain-predictive pattern

Given that we found shared neural representations between physical and affective vicarious pain (see Results for details), a general across physical and affective vicarious pain pattern was obtained by classifying vicarious pain (physical and affective) versus control stimuli and further evaluated by predicting physical vicarious pain versus physical control and affective vicarious pain versus affective control separately through 10-fold cross validation procedures. We next constructed 10,000 bootstrap sample sets to visualize the voxels that made the most reliable contribution to the classification and to decode the cognitive relevance of the classifier with the resultant Z map using the Neurosynth ^21^. Moreover, to compare the general vicarious pain pattern with the physical and affective vicarious patterns, we examined the similarities between this general vicarious pain pattern and the physical and affective vicarious pain patterns, respectively.

### Generalizability of the vicarious pain pattern

To test the functional relevance and generalizability of the empathic-induced neural pain pattern the unthresholded whole-brain pattern of the general across physical and affective vicarious pain was applied to determine the behavioral and neural responses during actual pain induction. To this end data from a previous study employing different levels of thermal pain induction during fMRI was used details provided in refs. ^39,40^. Briefly, the dataset included n = 33 healthy, right-handed participants (22 females; mean ± SD age = 27.9 ± 9.0 years). Six levels of temperature (ranging from 44.3 to 49.3°C in increments of 1°C) were administered during fMRI acquisition. During each trial, thermal stimulations were delivered to the left volar forearm alternating between runs. Each trial consisted of a 12.5s stimulus (3s ramp-up and 2s ramp-down periods and 7.5s at the target temperature), a jittered 4.5-8.5 sec delay, a 4 sec painful/non-painful decision period, and a 7s continuous warmth or pain rating period (on a visual analogue scale). In the first-level GLM analysis contrasts of interest included the 6 thermal pain stimulation levels. The general vicarious pain pattern from the current study was used to estimate the pattern expressions of each participant in each condition and next the neural pattern expressions of the 6 pain levels were (1) correlated with the temperature levels (1-6) as well as the subjective pain ratings separately and, (2) employed to discriminate high thermal pain stimulation (average of 48.3 and 49.3°C) versus low stimulation (average of 44.3 and 45.3°C), high stimulation versus medium stimulation (average of 46.3 and 47.3°C), as well as medium stimulation versus low stimulation. Moreover, we conducted the same analyses with physical and affective vicarious pain patterns to determine the robustness of the prediction (see **Supplementary results**).

## Data availability

Statistical and pattern weight images are available on Neurovault (https://neurovault.org/collections/6332/). Other data can be obtained from the corresponding authors upon request.

## Code availability

Code is available from the corresponding authors upon request.

## Acknowledgements

This work was supported by the National Key Research and Development Program of China (Grant No. 2018YFA0701400), National Natural Science Foundation of China (NSFC, No 91632117, 31700998, 31530032); Fundamental Research Funds for Central Universities (ZYGX2015Z002), Science, Innovation and Technology Department of the Sichuan Province (2018JY0001); National Institute of Mental Health (R01 MH116026) and National Institute of Biomedical Imaging and Bioengineering (R01EB026549).

## Author contributions

F.Z. and B.B. conceptualized the experiment, analyzed the data and wrote the original draft. J.L., L.X., X.Z. and M.F. acquired the data. J.L., W.Z., S.Y., K.M.K. and T.D.W. edited and revised the paper for important intellectual content.

