## Supplementary_Information for "Emotional contagion of pain across different social cues shares common and process-specific neural representations"

**Zhou et al.**

**Supplementary results**

**Generalization of pain-predictive patterns**

To test the generalizability of the identified neural patterns across different types of pain, we applied whole-brain physical and affective vicarious pain-predictive patterns to actual pain data using dot-product of vectorized activation maps with the pattern classifier weights separately. We found that both physical and affective vicarious pain-predictive pattern expressions were significantly correlated with subjective pain ratings (physical: $r_{196} = 0.417$, $p < 0.001$; affective: $r_{196} = 0.412$, $p < 0.001$) as well as pain intensities (physical: $r_{196} = 0.493$, $p < 0.001$; affective: $r_{196} = 0.412$, $p < 0.001$). Moreover, the physical vicarious pain-predictive pattern discriminated high thermal pain versus low thermal pain with a 97% accuracy (± 3.0% SE, $p < 0.001$, d = 2.12), high thermal pain versus medium thermal pain with a 88% accuracy (± 5.7% SE, $p < 0.001$, d = 1.54) and medium thermal pain versus low thermal pain with a 79% accuracy (± 7.1% SE, $p = 0.001$, d = 1.54) using a two-alternative forced choice test; affective vicarious pain-predictive pattern could also classify above chance for high thermal pain versus low thermal pain (85% ± 6.2% SE, $p < 0.001$, d = 1.11), high thermal pain versus medium thermal pain (85% ± 6.2% SE, $p < 0.001$, d = 0.88) and medium thermal pain versus low thermal pain (85% ± 6.2% SE, $p < 0.001$, d = 0.94) (**Supplementary Fig. 1A,B**).

When predicted with mid-insula partial patterns, physical pain-predictive pattern could discriminate high thermal pain versus low thermal pain (accuracy = 85% ± 6.2% SE, $p < 0.001$, d = 1.57), high thermal pain versus medium thermal pain (accuracy = 85% ± 6.2% SE, $p < 0.001$, d = 1.31) and medium thermal pain versus low thermal pain (accuracies = 82% ± 6.7% SE, $p < 0.001$, d = 1.38) above chance level; affective pain-predictive pattern could also correctly classify for high thermal pain versus low thermal pain (accuracy = 94% ± 4.2% SE, $p < 0.001$, d = 1.71), high thermal pain versus medium thermal pain (accuracy = 85% ± 6.2% SE, $p < 0.001$, d = 1.29) and medium thermal pain versus low thermal pain (accuracies = 91% ± 5.0% SE, $p < 0.001$, d = 1.60) (**Supplementary Fig. 1C,D**). In addition, both insula partial pain-predictive pattern expressions highly correlated with individual pain ratings (physical: $r_{196} = 0.451$, $p < 0.001$; affective: $r_{196} = 0.467$, $p < 0.001$) as well as thermal pain intensities (physical: $r_{196} = 0.462$, $p < 0.001$; affective: $r_{196} = 0.478$, $p < 0.001$).

**Supplementary Fig. 1.** Generalizability of the physical and affective vicarious pain-predictive patterns. Both whole-brain (A) and mid-insula (B) vicarious pain-predictive patterns could accurately predict the severity and classify the levels of self-experienced pain in an independent dataset.

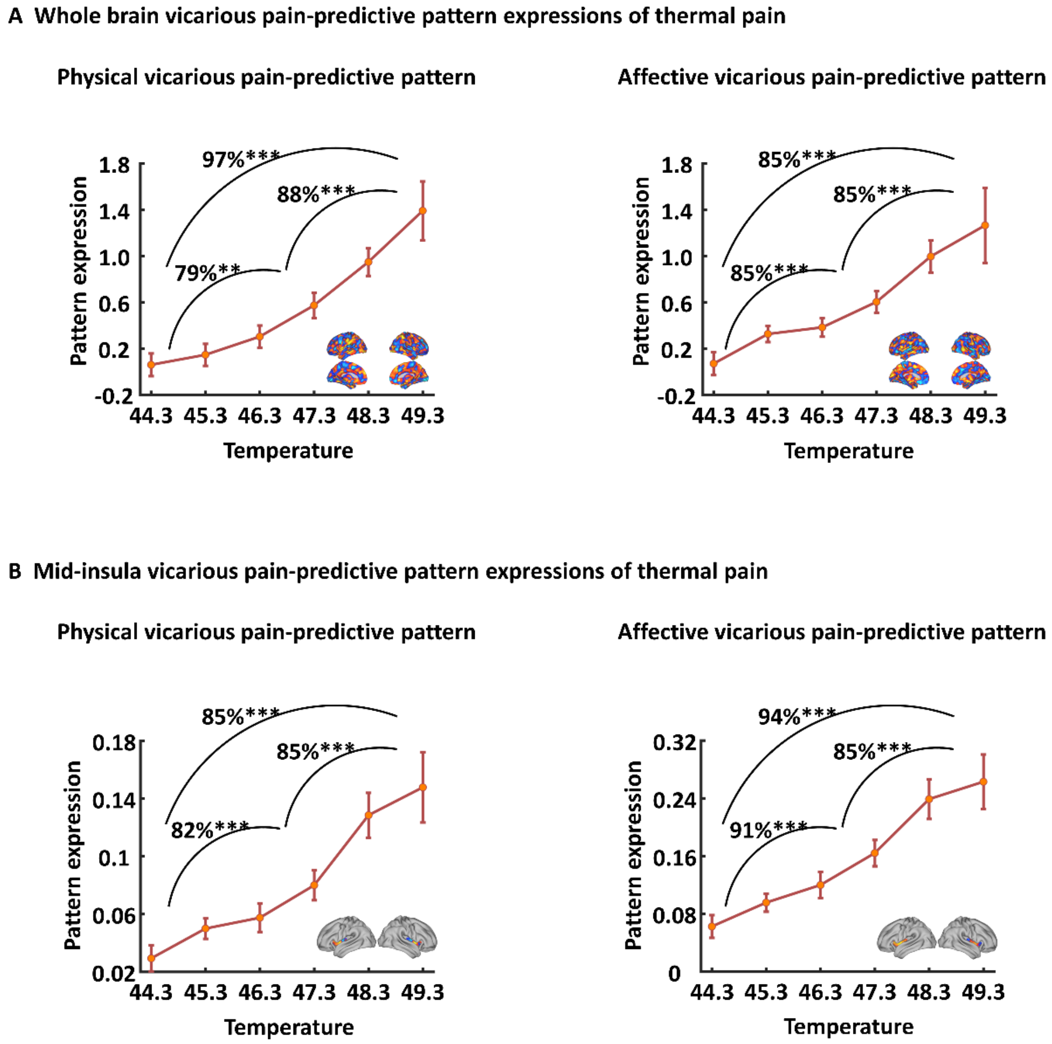

**Supplementary Table 1. Subjective ratings for stimuli (*M* ± *SD*)**

| Ratings | Categories of stimuli | | | |
|---|---|---|---|---|
| | affective control | physical control | Affective vicarious pain | physical vicarious pain |
| Pain intensity | 5.09 ± 7.34 | 8.49 ± 9.68 | 53.64 ± 21.43 | 71.43 ± 17.18 |
| Arousal | 16.79 ± 18.72 | 22.23 ± 18.74 | 54.48 ± 20.13 | 72.57 ± 17.53 |
